# Unlocking the biological potential of *Euglena gracilis*: evolution, cell biology and significance to parasitism

**DOI:** 10.1101/228015

**Authors:** ThankGod E. Ebenezer, Martin Zoltner, Alana Burrel, Anna Nenarokova, Anna M. G. Novák Vanclová, Binod Prasad, Petr Soukal, Carlos Santana-Molina, Ellis O’Neill, Nerissa N. Nankissoor, Nithya Vadakedath, Viktor Daiker, Samson Obado, Andrew P. Jackson, Damien Devos, Julius Lukeš, Michael Lebert, Sue Vaughan, Vladimir Hampl, Mark Carrington, Michael L. Ginger, Joel B. Dacks, Steven Kelly, Mark C. Field

## Abstract

Photosynthetic euglenids are major components of aquatic ecosystems and relatives of trypanosomes. *Euglena gracilis* has considerable biotechnological potential and great adaptability, but exploitation remains hampered by the absence of a comprehensive gene catalogue. We address this by genome, RNA and protein sequencing: the *E. gracilis* genome is >2Gb, with 36,526 predicted proteins. Large lineage-specific paralog families are present, with evidence for flexibility in environmental monitoring, divergent mechanisms for metabolic control, and novel solutions for adaptation to extreme environments. Contributions from photosynthetic eukaryotes to the nuclear genome, consistent with the shopping bag model are found, together with transitions between kinetoplastid and canonical systems. Control of protein expression is almost exclusively post-transcriptional. These data are a major advance in understanding the nuclear genomes of euglenids and provide a platform for investigating the contributions of *E. gracilis* and its relatives to the biosphere.

## Introduction

*Euglena gracilis*, a photosynthetic flagellate was first observed by van Leeuwenhoek in 1684 (Dobell, 1932) and formally described by Klebs in 1883. There are over 250 known species in the genus *Euglena*, with ~20 predominantly cosmopolitan taxa, including *E. gracilis* (Kim, *et al.,* 1998; Gojdics, 1953; Zakrys, 1988; Zakrys and Walne, 1994). Euglena are facultative mixotrophs in aquatic environments (Buetow, 1982) and many possess a green secondary plastid (McFadden, 2001). Amongst many notable features of *Euglena* are the pellicle, an unusual proteinaceous structure, the cell surface (Drago, *et al.,* 1997) and a red eyespot, a defining feature amongst species of photosynthetic euglenids, and which facilitates phototaxis (Daiker, *et al.,* 2011, van der Horst and Hellingwerf, 2004, Heijde and Ulm 2012, Iseki et al. 2002, Masuda, et al., 2013, Richter, et al., 2007). Euglenids belong to the Excavata supergroup, and thus are related to the Kinetoplastids and the Diplonemids. The former includes *Trypanosoma* and *Leishmania* (Adl, *et al.,* 2012), important unicellular parasites; the latter are little studied yet represent one of the most diverse and abundant lineages in the ocean. *E. gracilis* is therefore an important organism due to its evolutionary history, divergent cellular architecture and complex biology, together with a considerable potential for biotechnological exploitation (O’Neill, *et al.,* 2015). However, the full complexity of euglenid biology remains to be revealed, and the absence of a genome sequence or complete transcriptome has greatly hampered efforts to understand these organisms or to develop genetic tools (O'Neill, *et al.,* 2015).

Comparisons between free-living bodonid and euglenid relatives, early-branching trypanosomatids such as *Paratrypanosoma confusum* and fully parasitic forms, has provided essential information on the genetic changes associated with parasitism (Lukes, *et al.*, 2014, Flegontov, *et al.,* 2013, Jackson 2016). Both the cell surface and flagellum are of significant importance to life cycle progression, interactions with the environment and, in the case of parasitic trypanosomes, pathogenesis and immune evasion (Langousis and Hill, 2014, Perdomo, *et al.,* 2016). The surface macromolecules of trypanosomatids are highly lineage-specific with roles in life cycle progression (Jackson et al., 2013, 2016, Kalb, *et al.,* 2016, Zoltner et al., 2016, Hovel-Miner et al., 2016, Devault and Banuls, 2008, Chamakh-Ayari et al., 2014), but it remains to be determined to what extent *E. gracilis* shares surface protein families or other aspects of their cell biology with parasitic kinetoplastids. Such information is valuable to understanding how parasitism arose.

*E. gracilis* produces a wide range of secondary metabolites, many of which are of potential commercial value (O'Neill, *et al.,* 2015). Euglena species are extremely resistant to conditions such as low pH and high metal ion concentrations, fuelling interest in possible utility as sentinel species and/or bioremediation agents (Yoshida, *et al.,* 2016, Miazek, *et al.,* 2015, Rodríguez-Zavala, *et al.,* 2007, dos Santos Ferreira, *et al.,* 2007). *E. gracilis* is also used as a food supplement, particularly in Japan. Furthermore, the biochemical complexity of *E. gracilis* is of considerable potential value for biotechnology and for biofuel production (Mahapatra, *et al.,* 2013; Furuhashi et al., 2015; Yamada et al., 2016).

*E. gracilis* possesses a complex genome, organized into nuclear, chloroplast and mitochondrial elements. The coding potential of the mitochondrial genome is surprisingly small (Dobáková, *et al.,* 2015, Faktorová, *et al.,* 2016), whilst the chloroplast genome is more conventional (Hallick, *et al.,* 1993). The nuclear genome size of *E. gracilis* has been estimated as between one and nine gigabases (Dooijes et al., 2000; Stankeiwicz et al., 1983; Marzus et al., 1984, Ebenezer et al., 2017), and the organization and intron/exon boundaries for only a few genes have been described (Schantz and Schantz 1989; Jackson et al. 2006; Levasseur et al. 1994; Milanowski et al. 2014, 2016; Canaday et al. 2001). *E. gracilis* can support multiple splicing pathways, including both conventional and non-conventional *cis*-splicing (Milanowski *et al.,* 2014, 2016), as well as *trans*-splicing (Tessier, *et al.,* 1991). This level of complexity indicates that control elements for expression, transcription and mRNA maturation pathways remain essentially uncharacterised. Moreover, the interplay between genes specific to *E. gracilis*, those shared with kinetoplastids and/ or other lineages and those acquired during secondary endosymbiosis are unknown.

While the organellar genomes of Euglena have been sequenced, the nuclear genome remains uncharacterised. To address this we undertook genomic, transcriptomic and proteomic analysis of the Z1 strain of *E. gracilis* to access multiple aspects of euglenid biology.

## Results

### A sequence resource for *Euglena gracilis* reveals an extensive predicted proteome but minuscule coding content

For sequencing genomic DNA, multiple draft assemblies were produced and the version with highest utility contained 2066288 scaffolds and contigs with N_50_ of 955. This assembly was selected on the basis that, although it was highly fragmented, the largest proportion (ribosomal RNA filtered) of total RNAseq reads could be mapped to this assembly (87%). Together, these data correspond to a total of 1.4Gbp and an estimated nuclear genome size of ~2Gbp at ~300x coverage. Based on CEGMA, we estimate that the genome is ~20% complete in terms of open reading frames (ORFs) and hence can only support an initial analysis of genome structure (Table S1).

None-the-less, restricting analysis to larger contigs >10kb did allow identification of several highly unusual aspects of *E. gracilis* genomic structure (Table 1, Figure 1 and S1). Out of 2066288 assembled scaffolds and contigs only 1459 were >10kb. BLAST searching these sequences with the *de novo* assembled transcripts identified 108 that were taken forward for gene structure analysis (query = euglena transcriptome; target = >10kb euglena genome; e value <= 0.001). Out of these 108 sequences with significant BLAST hit, full length assembled transcript alignments using Exonerate could be generated for 53. This contig cohort encompasses about 22Mb of sequence, but only 135 genes, suggesting an extremely low gene density. In the few contigs with coding sequence, there is frequently more than one ORF, suggesting ORF clusters within large expanses of non-coding sequence. Most recovered genes have evidence for cis-splicing, most such introns are conventional with a smaller proportion of intermediate and non-conventional splice sites (Breckenridge et al., 1999). Furthermore, some genes are apparently unspliced (Figure 1; Transcript 109) and in at least one case there is evidence that the gene may be subject to alternate splicing (Figure 1; Transcripts 524, 326, and 454). Thus, *E. gracilis* possesses an open reading frame (ORF) structure that is similar to many other lineages in terms of cis-splicing, but which is less dense and more complex than previously appreciated. The near complete absence of cis-splicing from bodonids and trypanosomatids reflects intron loss at the root of the kinetoplastidea (Mair et al., 2000).

**Table 1.** Statistics for contigs assembled and spanning greater than 10kb.

| **Contigs** | Total contigs analysed > 10 kb | **1459** |
| --- | --- | --- |
|  | Total nucs in contigs analysed | 22Mb |
|  | Contigs with CDS | 53 |
|  | Percent contigs with CDS | 3.6 |
| **CDS** | Number analysed | 135 |
|  | Average length | 3790 |
|  | Total length | 481369 |
| **Exons** | Number of exons analysed | 421 |
|  | Average Length | 174.54 |
|  | Median Length | 112 |
|  | Total Length | 73482 |
|  | Average per predicted CDS | 3.85 |
| **Introns** | Total introns analysed | 271 |
|  | Average Length | 1027.14 |
|  | Median Length | 598 |
|  | Total Length | 278354 |
|  | Introns per predicted CDS | 2.01 |
|  | Number/percent conventional | 218/80.1 |
|  | Number/percent intermediate | 30/11.1 |
|  | Number/percent non-conventional | 23/8.5 |
|  | Percent nucleotides in CDS (exon) | **0.003** |

**Figure 1.**
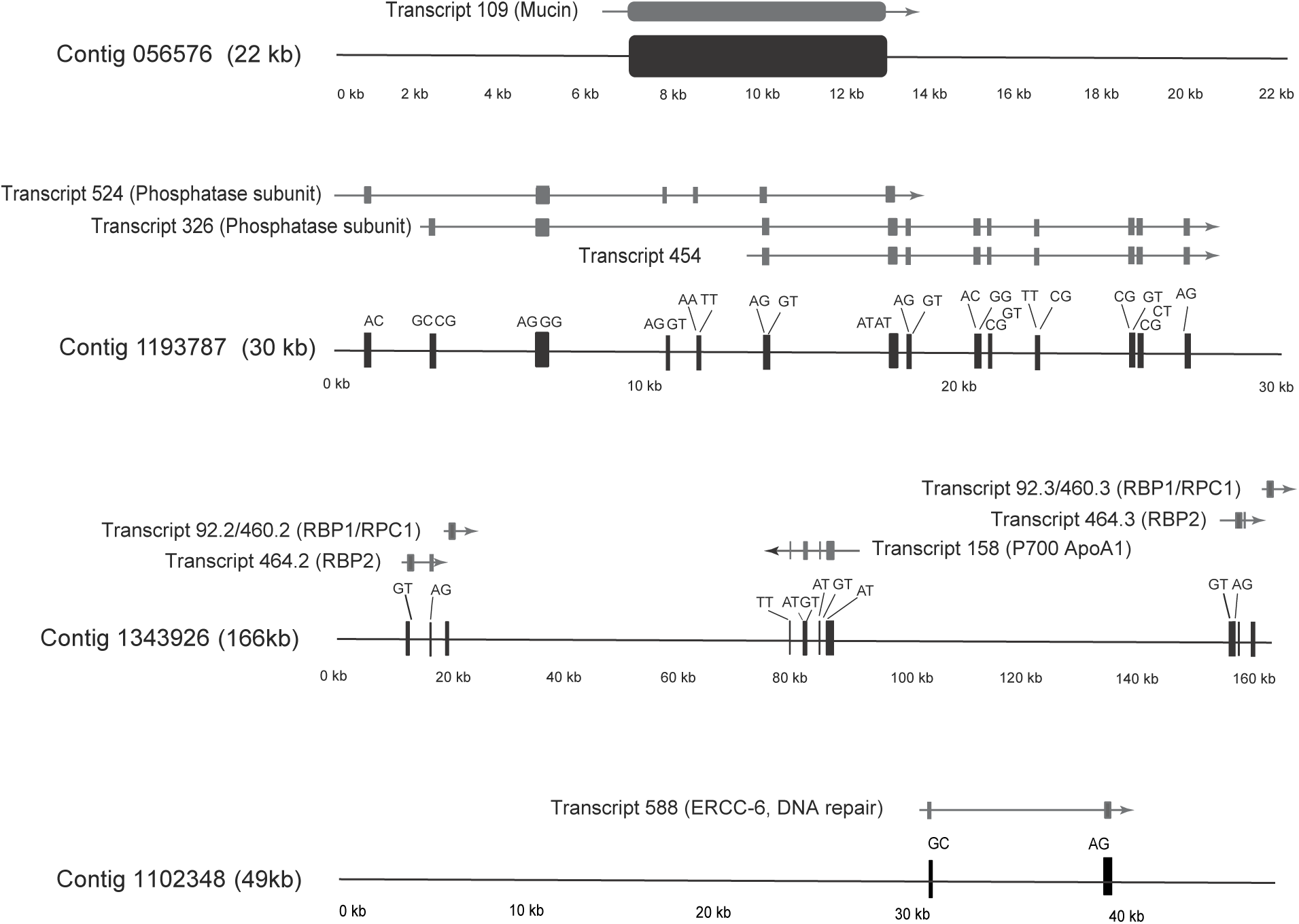
*Euglena gracilis* exonic structure. The predicted gene structure of several selected contigs is shown, including the mapped transcripts, predicted splice sites and intergenic regions. Note that transcripts 524 and 326, which encompass essentially the same portions of the genome, demonstrate possible differential exon inclusion, indicating differential open reading frame organisation and possible alternate splicing. Black boxes indicate exons, with predicted splice site dinucleotides indicated above. Transcripts are shown as arrows with the arrowhead indicating the predicted direction of transcription. Protein product annotations are indicated in parentheses. Contig sizes are shown in kb, and note that each contig is not drawn to the same scale. Further examples of predicted contig gene organisation are given in Figure S1.

Coverage by the transcriptome was significantly superior to the genome. Here CEGMA analysis indicated the presence of 87.9% conserved genes, which compares very favourably to *T. brucei* at 82.66% (Table S1D). The total number of *E. gracilis* ORFs (36 526) is two to four times greater than *Bodo saltans* (18 963), *T. brucei* (9068), *T. cruzi* (8272) or *Naegleria gruberi* (15727) (Jackson, *et al.,* 2008, Berriman, *et al.,* 2005, Ivens, *et al.,* 2005, Fritz-Laylin, *et al.,* 2010). Blast_2_Go identified 16 103 sequences with GO terms (Figure S2) and InterProScan increased the number of annotated genes by ~20%. Significantly, comparisons between the genome and transcriptome indicates that very little of the genome is coding sequence, with the contigs containing a vanishing 0.003%. Overall, these assemblies reconstruct a highly unusual eukaryotic genome organization while still providing data of sufficient quality for robust annotation and prediction of function, as well as for proteomic analyses.

### Post-transcriptional control of protein expression is similar to trypanosomatids

To improve annotation and investigate the control of gene expression, we conducted a label free comparative proteomic analysis between light and dark-adapted *E. gracilis*, as under these well-controlled conditions significant changes to the proteome were expected. Following a period of dark-adaptation, we confirmed by UV/VIS spectroscopy and SDS-PAGE that photosynthetic pigments were lost and that ensuing ultrastructural changes were as expected (Figure S3). Total protein extracts were separated by SDS-PAGE and 8661 distinct protein groups were identified by LCMS. Ratios for expression levels between both light and dark conditions could be determined for 4681 protein groups (Table S2), including a cohort of 384 protein groups that were observed in only one state (232 in light and 152 in dark). Protein groups not identified at the peptide level in all replicates for one state were rejected from further analysis, but are included in Table S2. In parallel, we extracted RNA for RNAseq analysis and comparing the 41045 transcripts identified from this dataset with the protein groups provided a set of 4287 genes with robust data for both RNA and protein.

The correlation between transcript and protein abundance changes was extremely poor (Figure 2, S2). While consistent with earlier studies (Van Assche, *et al.,* 2015, Araújo and Teixeira, 2011) these data extend the analysis to encompass several thousand proteins, and hence indicate that the absence of transcriptional control for nuclear genes is a general feature and not specific to a restricted gene cohort. Similarly, the majority of the 384 protein groups detected in only one condition were not reflected in changes in their RNA levels, i.e. transcripts were detected in both conditions with similar intensity. Transcripts with differential abundance that did correlate with protein abundance are encoded by the chloroplast genome, including several photosystem I proteins, including P_700_ chlorophyll apoprotein A_1_ and chloroplast encoded translation elongation factor EF-Tu. Nuclear translation elongation factors are not influenced by switching growth conditions from dark to light (Montandon et al., 1989), consistent with our finding of no differential expression of nuclear EF-1α, while both the chloroplast EF-Tu protein and corresponding transcript [EG_transcript_1495] are highly upregulated in the light. The absence of transcriptional control is highly similar to that reported for the kinetoplastids, despite the presence of widespread *cis*-splicing in *E. gracilis* and which indicates an absence of promotor-based modulation of transcriptional activity.

**Figure 2.**
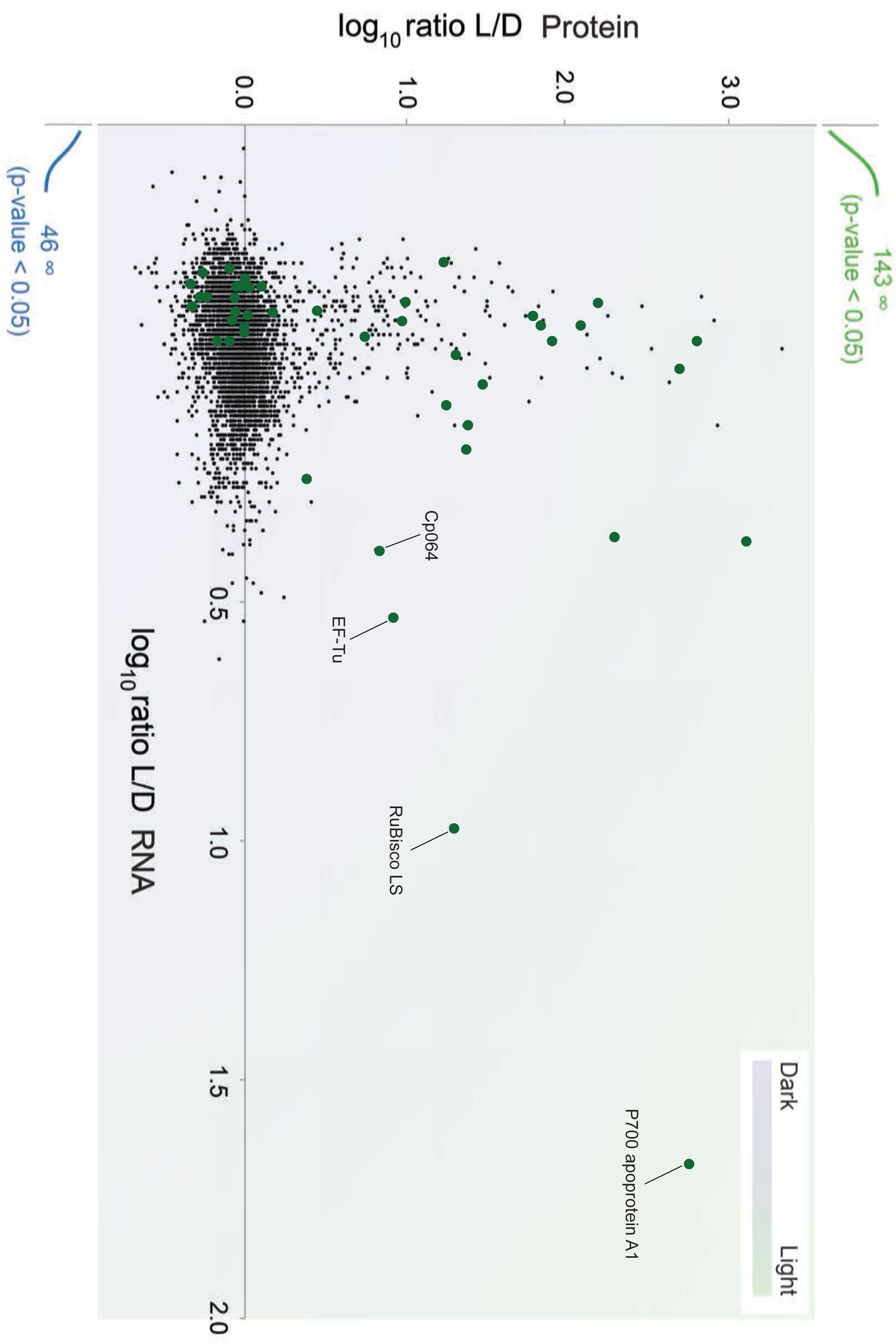
Control of expression level is mainly post-transcriptional in *E. gracilis.* Alterations to the transcriptome and proteome in response to ambient light or complete darkness were analysed using RNAseq and SILAC/LCMS^2^ proteomics respectively. Data are plotted for individual transcripts/polypeptides as the log_10_ ratio between the two conditions, light (L) and dark (D), with protein on the y-axis and RNA on the x-axis. The presence of a number of proteins that were detected exclusively under one or other condition (hence infinite ratio) are indicated in green (for light) and blue (for dark). With the exception of three transcripts, which are plastid encoded (labeled), there is little alteration to RNA abundance, but considerable changes to protein levels. Raw data and transcriptome/proteome analysis are provided in supplementary data.

### Extensive gene families from multiple evolutionary origins

To identify ortholog families and understand their evolutionary dynamics, we used Orthofinder (Figure 3 and S3). As expected, the phyletic pattern that contained the largest number of orthogroups was one that contained representation from all supergroups, and dominated by core metabolic, structural and informational processes. A second cohort are shared between *E. gracilis* and excavates. These classes are broadly within the relative frequencies of what was expected from previous analysis of excavate genomes (Simpson, *et al.,* 2006) and also with the phylogenetic positions of the relevant taxa (Figure 3, Table S3). A third cohort represents likely nuclear transfer of endosymbiotic genes as a result of acquisition of the plastid and mitochondrion. Consistent with these reports, genes that are shared only between *E. gracilis* and disparate photosynthetic lineages may represent lateral gene transfer events. To determine if other lateral gene transfer events could be detected in orthogroups that contained other non-photosynthetic lineages, each othrthogroup was subject to analysis using Notung. While multiple candidate lateral gene transfer events were detected, all originated from other photosynthetic lineages (Figure SX) and likely came from the green algal endosymbiont nucleus. GO terms associated with orthogroups indicated increased regulatory function genes in green/secondary plastid orthogroups, whilst phosphorus-dependent metabolic processes predominate among orthogroups shared with kinetoplastids (Figure S4, Table S4).

**Figure 3.**
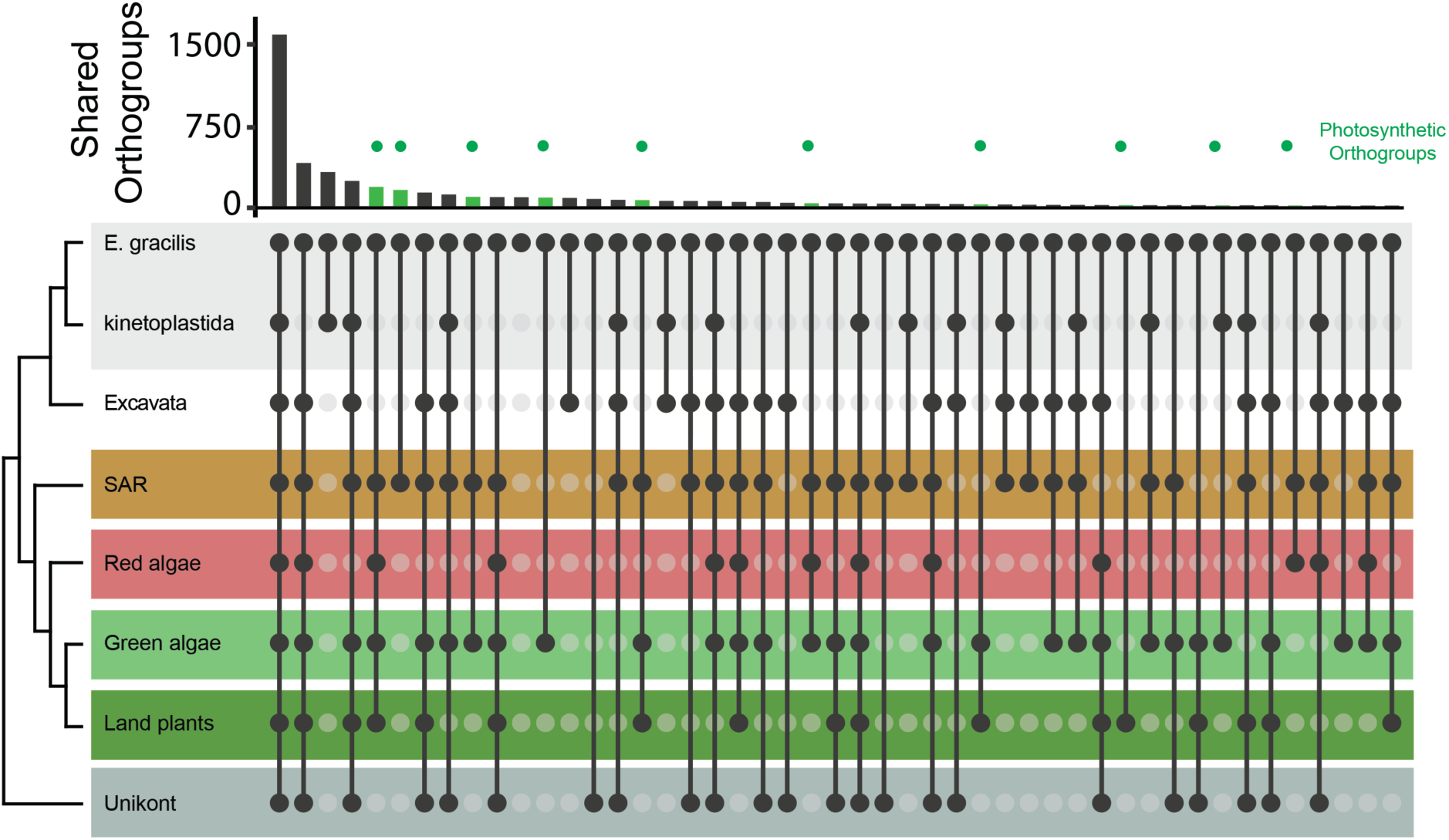
*Euglena gracilis* shares orthologs with a diverse array of lineages. Panel A (top): Histogram of *E. gracilis* orthologous groups clustering with selected eukaryotic lineages as determined with Orthofinder. The X-axis shows the number of orthogroups and y-axis shows the taxon groupings representative of selected eukaryotic groups. Histogram bars highlighted in green indicate orthogroups shared with photosynthetic organisms. Panel A (lower): Taxa sharing orthogroups with *E. gracilis,* where black circles correspond to the presence of orthogroup members while light grey circles correspond to the absence of orthogroup members in the genome. Black tie bars linking black circles are for clarity only. Eukaryotic taxon groupings are colored accordingly: grey, Euglena and kinetoplastida; white, other members of the Excavates excluding Euglenozoa; brown, SAR, pink, red algae; light green, green algae; dark green, land (vascular) plants and dark grey, Unikonts. An expanded version of this figure, broken down by species is given as Figure S2. Panel B: The number of shared orthogroups, binned by various taxon groupings.

We also performed exhaustive analysis to establish ancestry of all proteins from the predicted *Euglena* proteome using single protein phylogenies. These analyses should reveal genes with probable vertical ancestry and those more closely related to more distant organisms as evidence for gene transfer events. The latter may be regarded as products of genes acquired horizontally, although the direction of transfer cannot be objectively established and cases of gene transfers from *Euglena* may also be present in the data set. Out of 36,526 predicted genes only 4,087 formed a robust (bootstrap support >=75%) sister relationship with a taxonomically homogeneous clade and these were binned accordingly (Figure 3). Because we adopted relatively strict criteria, these numbers are conservative estimates. Out of the 4,087 proteins with robustly supported sister lineages, 1,816 (44%) were related to one of the lineages of Excavata present in the database and 1,420 (35%) were related specifically to kinetoplastids. This major fraction represents the vertical inherited component of the genome. Another large set speaks to the origin of the secondary plastid and will be treated below. Finally, the proteins in the bins of Prokaryota (220) and non-photosynthetis eukaryotes, e.g. Metazoa (149) and Amoebozoa (145), are most probably the results of horizontal gene transfers but could also be artefacts of our methodology. We also noted that several orthogroups contained a very large number of Euglena sequences (Figures S4 and 5; Tables S2-S3), and these were analysed phylogenetically and annotated for architecture (Figure 4 and S5).

A family of nucleotidylcylase III (NCIII)-domain-containing proteins (OG000541) that are widely distributed across eukaryotes are highly significant, in terms of possible functions and in size. In trypanosomes, adenylate cyclases are mediators of immune modulation in the mammalian host (Salmon, *et al.,* 2012). One nucleotidylcylase subfamily is restricted to kinetoplastids and organisms with secondary endosymbionts and contains photosensor adenylate cyclases described previously (Iseki et al, 2000). These proteins possess one or two BLUF domains (blue light sensor) with a double NCIII domain (Figure 4), and are phylogenetically similar to the NCIII-family of *N. gruberi* (Fritz-Laylin *et al.,* 2010). A second subfamily is pan-eukaryotic and possesses one NCIII domain, several *trans*-membrane domains plus HAMP and cache1 domains involved in signal transduction, particularly in chemotaxis (Dunin-Horkawicz and Lupas, 2010, Anantharaman and Aravind, 2000). Again, this subfamily is closely related to a group of proteins encoded by the *N. gruberi* genome. The third subfamily belongs to a kinetoplastid only cluster with trans-membrane proteins and frequently also HAMP and cache1 domains. The overall complexity of this family suggests very great flexibility in nucleotidylcyclase evolution and that many lineage-specific paralogs have arisen with implications for signal transduction, likely an important regulatory pathway in *E. gracilis*.

**Figure 4.**
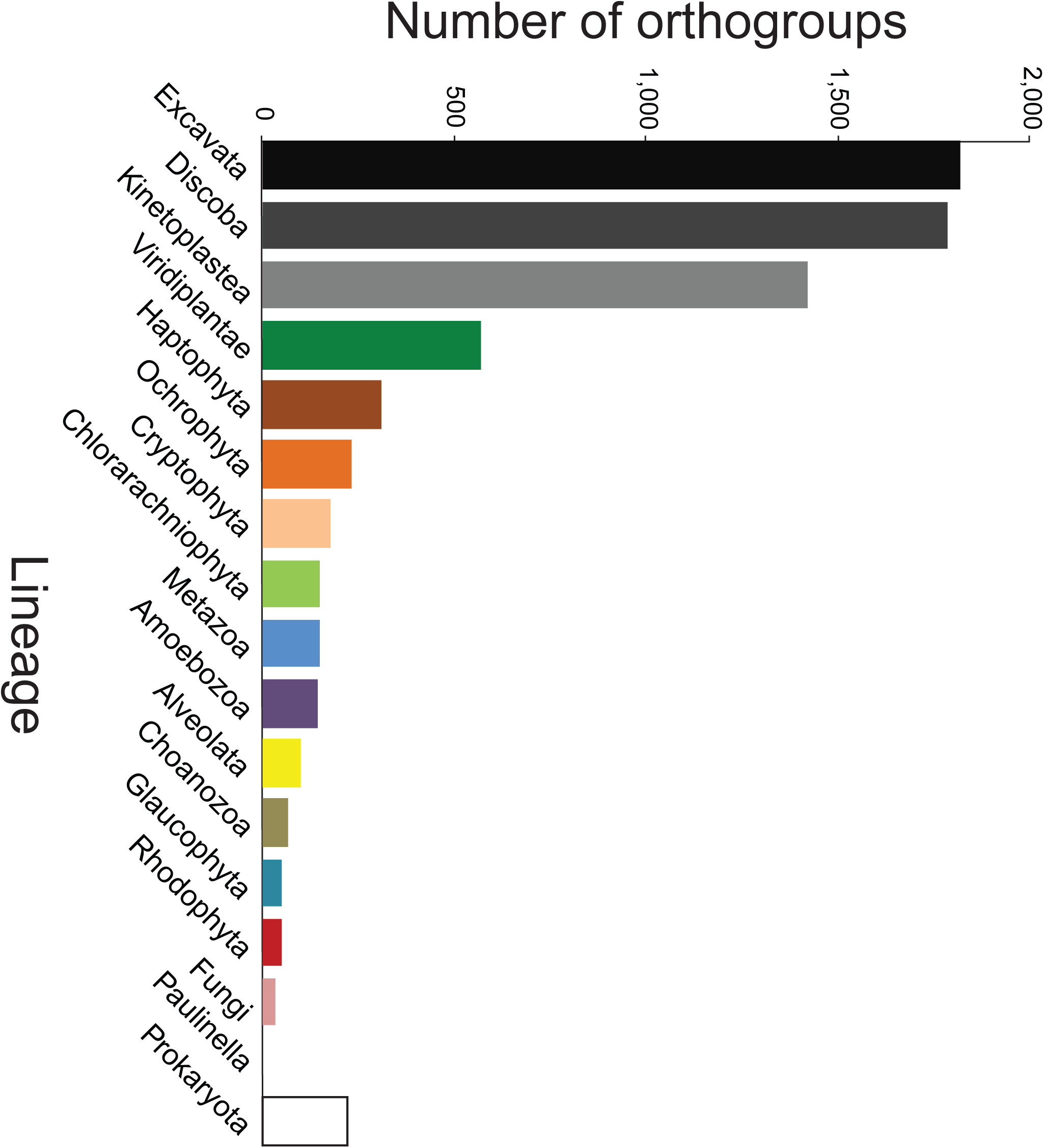
Large paralog gene families are present in the *Euglena gracilis* genome. Several orthogroups contain many Euglena paralogs. The phylogenetic distribution of one such large orthogroup (OG000541), the nucleotidylcyclase III domain-containing proteins, is shown. Lineage groupings are colour coded: Gray; all eukaryotes (and collapsed for clarity), Red; *N. gruberi*, Amber; *B. saltans* and Green; *E. gracilis*. Clades containing only Euglena sequences are boxed in green. Each sequence has been assigned a domain composition (color gradient black to teal to blue), number of predicted transmembrane domains (colour coded red to orange to black gradient). To obtain this phylogenetic tree, sequences with likely low coverage (less than 30% of the length of the overall alignment) were removed during alignment to avoid conflicting homology or artefact generation. Domain compositions identified are nucleotidylcyclase III, BLUF, NIT, P-loopNTPase, HAMP and Cache1. Additional examples are in Figure S5.

An extensive protein kinase C-domain-containing group of protein kinases was also recovered (OG000113 and OG000137), with extensive lineage-specific expansion in *E. gracilis* (Figure S5). A third orthogroup possesses a REC-domain (signal receiver domain); with clear lineage-specific *E. gracilis* paralogs (Figure S5). The *E. gracilis* members possess an H-ATPase domain, which is distinct from the PAS domain present in many of the predicted proteins from other lineages. The presence of families of signalling proteins that have expanded independently in *E. gracilis*, suggests both highly complex and divergent signalling pathways. These very large paralog cohorts likely partly explain the expanded coding potential, as well as contribute towards the ability of *E. gracilis* to respond to environmental changes.

#### Conservation and divergence between *E. gracilis* and kinetoplastids

The phylogenetic position of Euglena makes it an important organism for the better understanding of several critical features of the parasitic kinetoplastids. A unique feature of energy metabolism in kinetoplastids is compartmentalisation of the first six or seven glycolytic enzymes within peroxisome-derived glycosomes and the presence of additional enzymes used for metabolism of the glycolytic intermediate phospho-enolpyruvate to succinate (Szöör, *et al.,* 2014). Glycosomes have been recently detected in the other major euglenozoan group, the diplonemids (Morales et al., 2016). Using 159 query protein sequences of high confidence glycosomal *T. brucei* proteins (Güther et al. 2014), we found candidate orthologs for the majority, but no evidence that enzymes linked to carbohydrate metabolism were peroxisomal. Of 47 high confidence glycosomal proteins annotated as hypothetical or trypanosomatid-specific, none had a detectable ortholog in *E. gracilis*. This indicates that peroxisomes in *Euglena* are likely to serve primarily in diverse aspects of lipid metabolism, but that the evolutionary drive to kinetoplastid glycosomes, and possibly also in diplonemids, was accompanied by innovations extending beyond targeting of glycolytic enzymes to the glycosome.

The pellicle of *E. gracilis* is a remarkable striated proteinaceous structure in close association with a microtubule corset, with some similarity to the subpellicular array of trypanosomatids (Lonergan, 1985), but which is clearly structurally distinct. The plasma membrane composition of kinetoplastids is also lineage-specific, in terms of the major protein species present, and is a major contributor to host-parasite interactions, immune evasion and pathogenesis (Gadelha et al., 2015), while transporters and some additional surface protein families are conserved. To compare with *E. gracilis*, we predicted membrane proteins using the signal peptide together with orthogroup clustering. Many have significant similarity to kinetoplastid proteins (1103), *B. saltans* (32) or non-kinetoplastida (487) (Table S5). About 698 proteins with a signal peptide appear to be Euglena specific. Most *Euglena*-specific predicted membrane proteins are single-copy (87.5%), while there are clear large families that possess conserved features, as discussed above. Notably, we were unable to identify a rhodopsin homolog, which is in contrast to several biochemical analyses that have suggested the presence of retinal, the rhodopsin cofactor, and which have been interpreted as evidence for a rhodopsin-like light sensing transduction mechanism (Barsanti et al., 2009).

In common with *B. saltans, E. gracilis* has a distinct class of amastin, a major surface protein of the kinetoplastids. This arose from a single ancestral form shared with the last common ancestor of euglenozoans (Figure S6). *E. gracilis* also possesses enzymes required for synthesis of lipophosphoglycan (LPG), a conserved glycoconjugate in *Leishmania* and *Phytomonas,* implicated in defence and disease mechanisms, together with the GPI-anchor pathway. These data suggest that LPG predates the evolution of parasitism, and that the ancestral role was likely one of defence, possibly against proteases and predation, or could even have a role in cell-cell/cell-substrate interaction. Significantly, gp63, a major surface protein present in the vast majority of eukaryotes and involved in *Leishmania* pathogenesis is absent, and hence must represent a secondary loss following separation from the kinetoplastid lineage.

The endomembrane system is responsible for the biosynthesis, degradation and targeting of proteins and lipids. The complexity of the endomembrane compartments in trypanosomatids exhibits an evolutionary history connected with pathogenesis and life style and compartmental complexity is decreased in the trypanosomatids compared with free-living bodonids, and even further in African trypanosomes (Jackson et a., 2016, Venkatesh et al., 2017). *E. gracilis* possesses a relatively complete set of membrane-trafficking proteins, extending this trend further (Figure S7, Table S6). Two key complexes involved in vesicle coat formation, AP5 and TSET, are absent from the kinetoplastids. While AP5 is also absent from *E. gracilis* but a near complete TSET is present, indicating that loss of AP5 took place after the split between Heterolobosea and Euglenozoa but that loss of TSET was later. Significantly, endosomal pathways are more complex than kinetoplastids, with multiple Rab7 (lysosome) and Rab11 (recycling endosomes) paralogs, together with ER-associated events, with five and three paralogs for Rab1 (ER exit) and Rab32 respectively. Some of these may be associated with the contractile vacuole, an endolysosomal organelle responsible for osmoregulation in many freshwater protists, as well as highly sophisticated sorting mechanisms associated with nutrition.

In kinetoplastids an unusual cytoskeletal element, the bilobe, plays a central role in Golgi, flagellar pocket collar and flagellum attachment zone biogenesis (Zhou, *et al.,* 2010). All of the structural proteins (MORN1, RRP1, BILBO1, Centrin-2 and Centrin-4) were found (Esson, *et al.,* 2012; Morriswood, *et al.,* 2013; Wang, *et al.,* 2012; Zhou, *et al.,* 2010; McAllaster, *et al.,* 2015; Aslett, *et al.,* 2010; Bugreev, *et al.,* 2011). Therefore, the potential for the synthesis of a bilobe-like structure in *E. gracilis* is possible, although clearly experimental evidence is needed for the presence of such a structure (Gheiratmand and He, 2014).

The extreme size of the *E. gracilis* genome and complex splicing machinery suggests sophisticated mechanisms for organizing chromatin, mRNA processing and transcription are present (Breckenridge, *et al.,* 1999, Milanowski, *et al.,* 2016), and are likely more complex than kinetoplastids, as both *trans-* and several types of *cis-*spicing are present (Günzl, 2010, Michaeli, 2011). Furthermore, the *E. gracilis* nucleus has unusual heterochromatin organization, where electron-dense regions appear as numerous foci throughout the nucleoplasm (Figure S8). Several novel aspects of nuclear organisation are now known in kinetoplastida, including the lamina, kinetochores and some aspects of the nuclear pore complex.

Neither the widespread lamins nor the kinetoplastid-specific NUP-1/2 lamina are found in *E. gracilis* which indicates that chromatin anchoring to the nuclear envelope exploits a distinct mechanism from either kinetoplastids, metazoan or amoeba (Kořený and Field 2016). Much of the nuclear pore complex (NPC) is well conserved across evolution, but Euglena possesses orthologs of the trypanosomatid nuclear basket proteins Nup92 and Nup110, that replace Mlp1/2 of fungi and animals (Holden et al., 2014). Significantly, DBP5 and Gle1, two proteins involved in mRNA export in a broad range of taxa, with the exception of trypanosomes, are present in Euglena. This is consistent with an earlier hypothesis that the absence of Gle1 and Dbp5 is connected with the absence of *cis*-splicing (Obado et al., 2016). Together these data indicate the presence of a hybrid NPC, with high conserved elements together with specific modulations associated with the trypanosomatids.

Kinetochores, required for engagement of chromosomes with the mitotic spindle, in trypanosomes possess essentially none of the factors present in mammals or fungi (Figure S8, Table S8) (Akiyoshi and Gull, 2014, D’Archivio et al.,2017). Of the trypanosomatid kinetochore proteins, only KKT19 and KKT10 are clearly present in *E. gracilis*. As these are a kinase and phosphatase respectively, it is possible that they are associated with non-kinetochore functions in Euglena. Further, very few canonical kinetochore proteins were found, suggesting divergence from the higher eukaryote configuration as well as the trypanosomatids. Overall, these observations suggest unique mechanisms operating in the *E. gracilis* nucleus, and which may reflect the transition between conventional kinetochores, lamins and nuclear pores into the more radical configuration present in kinetoplastids.

#### A *distinct mitochondria configuration*

Providing the group its namesake, Kinetoplastids contain a mitochondrial genome (the kinetoplast) with a unique structure, comprising 35-40 copies of a maxicircle and encoding several mitochondrial proteins, and several thousand minicircles encoding guide RNAs required for editing of maxicircle transcripts (Read et al., 2016; Faktorová, *et al.,* 2016). This structure is attached to the flagellum basal body *via* a complex cytoskeletal element, the tripartite attachment complex (TAC). We find no evidence for RNA-editing in *E. gracilis*, nor for the TAC, both of which are consistent with the presence of a mitochondrial genome composed of only short linear DNA molecules (encoding only seven different proteins) and a conventional mitochondrial mRNA transcription system (Dobáková, *et al.,* 2015). Specifically, only 14 of 51 proteins involved in RNA-editing in *T. brucei* (David et al., 2015) had reciprocal best BLAST hits, and only one predicted protein contained a mitochondrial targeting signal. No homologs to TAC proteins were found (Table S6).

The predicted *E. gracilis* mitochondrial proteome consists of 1093 proteins, divided into 16 KEGG categories (Figure 5 and S10). This includes the seven protein-coding genes encoded by the mitochondrial genome (Dobáková, *et al.,* 2015; Buhrman, *et al.,* 2013). The kinetoplastid mitochondrion possesses a non-canonical outer mitochondrial membrane translocase ATOM (archaic translocase of the outer membrane). In kinetoplastids, TOM40, a very conserved beta-barrel protein that forms the conducting pore, is highly divergent, and referred to as ATOM40 (Pusnik et al., 2011, 2012 Žárský et al., 2012). We identified homologues of two specific receptor subunits of ATOM, namely ATOM46 and ATOM69 (Mani et al., 2016), and two TOM40-like proteins. Both of the latter are very divergent, and could not be assigned unequivocally as TOM40 orthologs.

**Figure 5.**
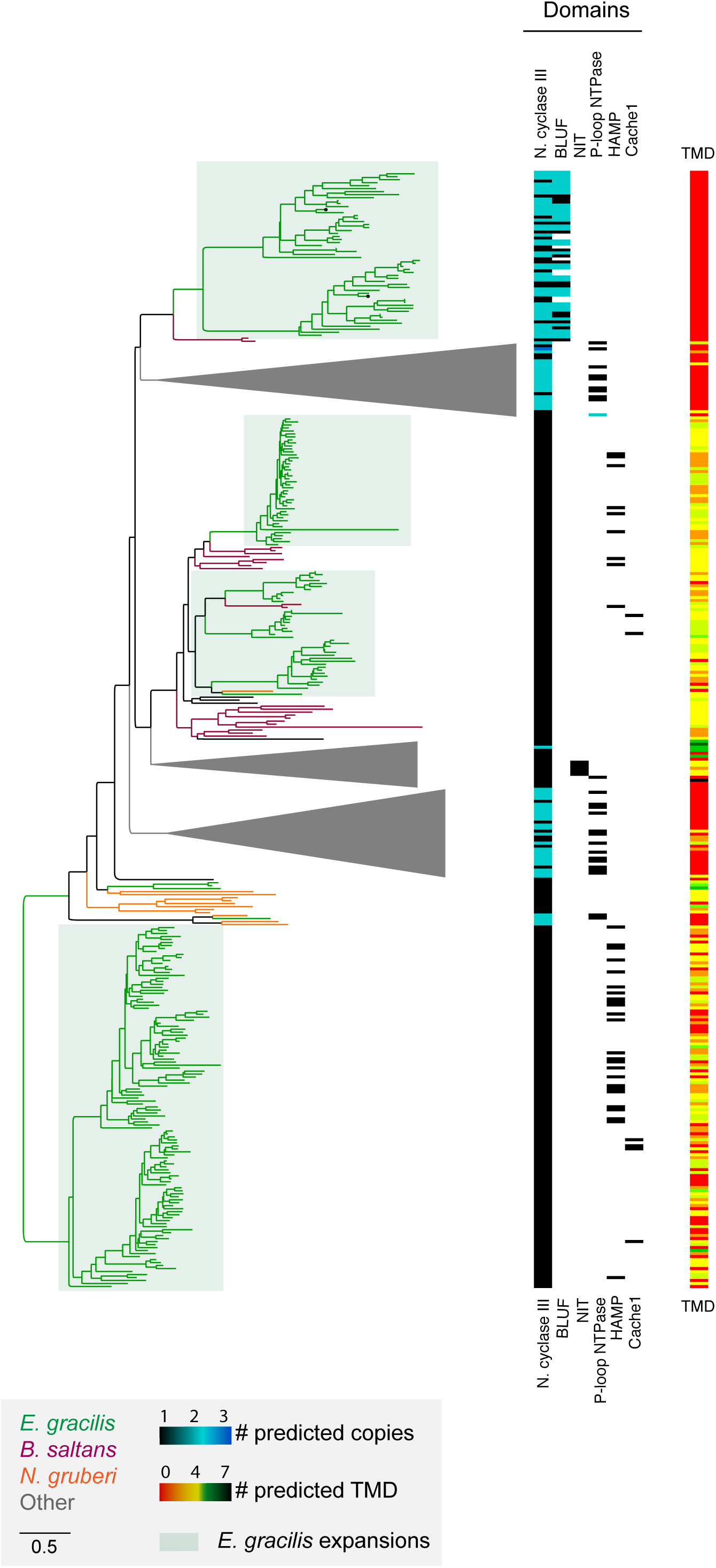
*Euglena gracilis* has flexible and fault-tolerant mitochondrial metabolism. Proteins involved in mitochondrial pathways and complexes are shown, including: tricarboxylic acid (TCA) cycle, pyruvate dehydrogenase, fatty acid metabolism, complexes I-V of respiratory chain, ubiquinone biosynthesis, sulfate assimilation pathway, Fe-S cluster assembly and export, TIM/TOM complex and mitochondrial import machinery. Colour codes: dark blue, nucleus encoded, present in predicted mitochondrial proteome; light blue, present in transcriptome without evidence for mitochondrial localization; light blue/white – mitochondrion-encoded proteins identified previously by Dobáková et al., 2015; grey – expected in nuclear transcriptome and not found; grey/white – expected in mitochondrial genome and not found. The *E. gracilis* mitochondrion can produce energy under both aerobic and anaerobic conditions and has workarounds for the main mitochondrial pathways, such as TCA cycle and respiratory chain, which may explain the outstanding adaptability of this organism.

We also identified canonical subunits of respiratory chain complexes I - V and 27 homologs of kinetoplastid-specific proteins, together with the widely represented alternative oxidase (TAO or AOX), consistent with earlier work (Perez et al., 2014). Moreover, an ortholog of *T. brucei* alternative type II NADH dehydrogenase (NDH2) was detected. We found only 38 of 133 canonical and only 3 of 56 kinetoplastid-specific mitoribosomal proteins, which suggests considerable divergence. Hence, the *E. gracilis* mitochondrion has unique features, representing an intermediate between the mitochondria familiar from yeast or mammals and the atypical biology of the organelle in kinetoplastids.

#### The plastid of Euglena

The Euglena plastid is a recent secondary acquisition, and represents a near unique case for studying fundamental aspects of both autogenous and endogenous organelle evolution. The predicted *E. gracilis* plastid proteome contains 1902 proteins (Figure 6, Table S6). Metabolic pathways and enzymes typical for the plastid are present, including 70 proteins involved in chloroplast electron transport chains and light harvesting antennae. A few undetected genes, such as glucose-6-phosphate isomerase in glycolysis/gluconeogenesis or 15-*cis*-phytoene desaturase in carotenoid biosynthesis pathway likely arise from incomplete sequence data, since both pathways are known in *Euglena* (Hallick et al., 1993). The C_5_ tetrapyrrole pathway was completely reconstructed and the C_4_ pathway for aminolevulinate synthesis is absent from the plastid, consistent with previous findings (Gomez-Silva et al., 1985). Enzymes connecting the cytosolic/mitochondrial mevalonate and plastidal methyl-d-erythritol pathway (MEP/DOXP) pathways of terpenoid synthesis were not found, in accordance with findings suggesting two isolated pools of geranylgeranyl pyrophosphate in plastid and in the cytosol of Euglena. Biosynthesis of carotenoids and non-plastidic isoprenoids (e.g. sterols, dolichols) is not connected (Kim et al., 2004). It should be noted that more than 50% of the predicted plastid proteome represent proteins with no significant homolog in the databases.

**Figure 6.**
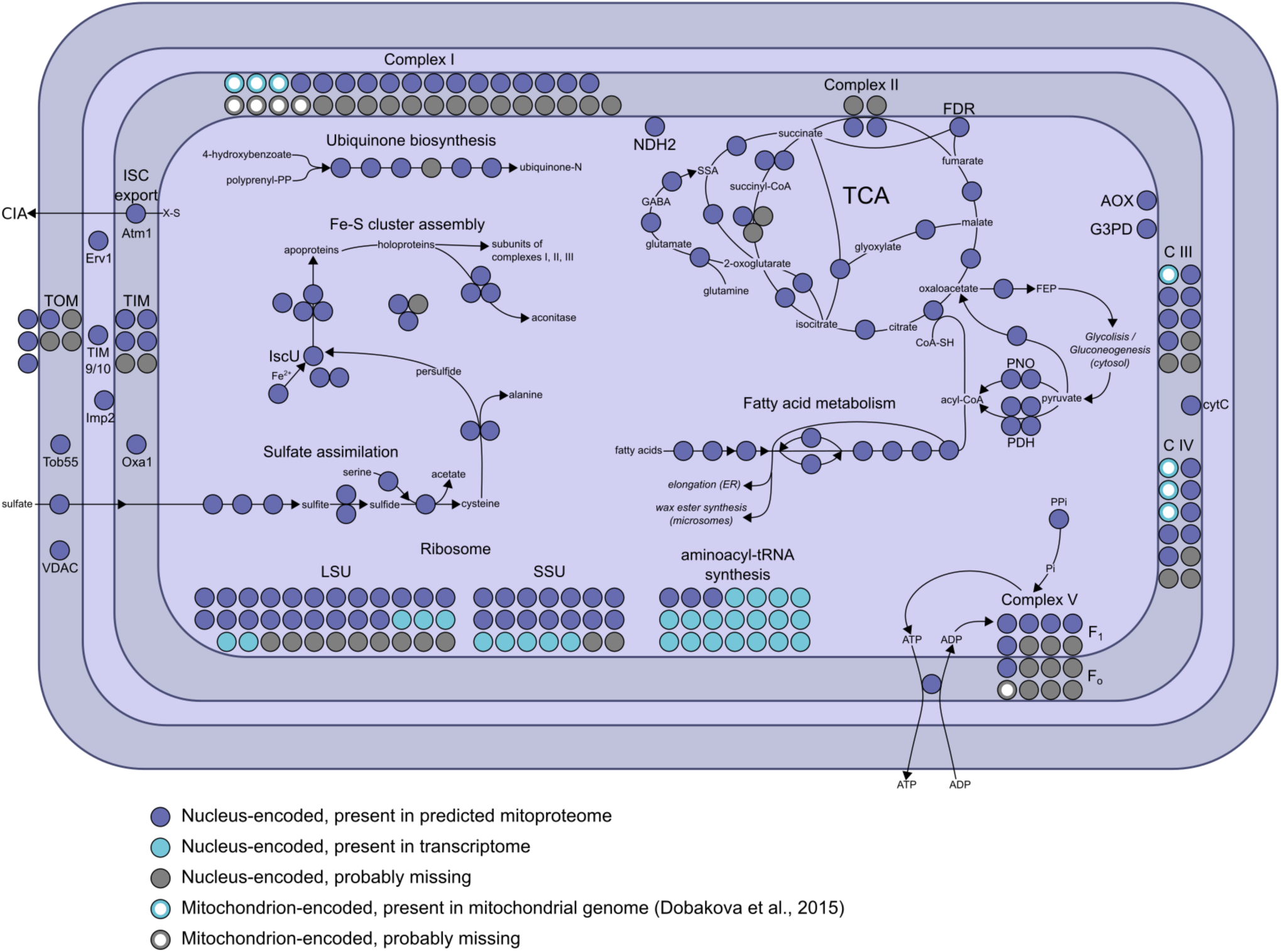
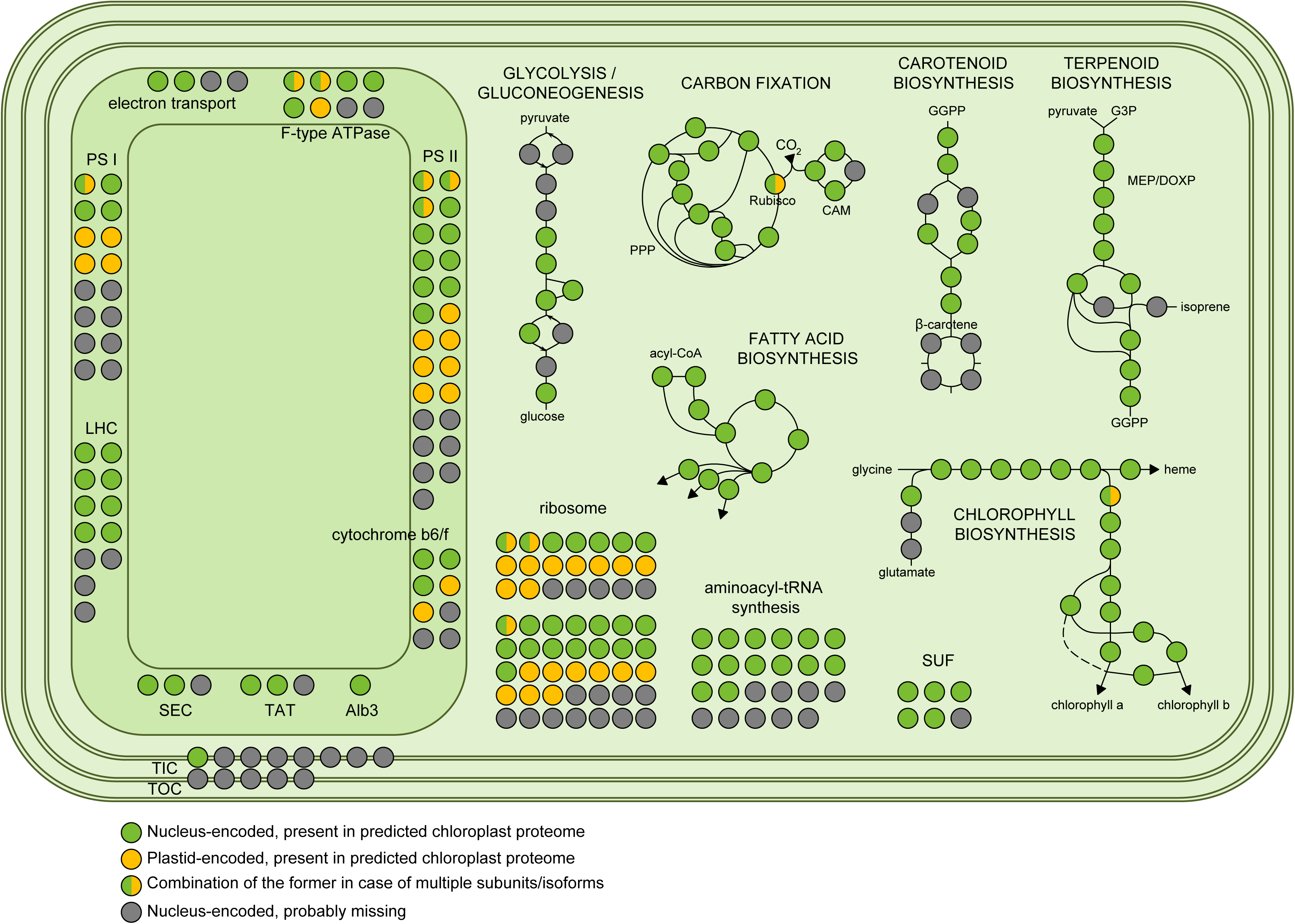
The *Euglena gracilis* plastid possesses broad metabolic potential. Proteins involved in core plastid metabolic pathways were identified and include: glycolysis/gluconeogenesis, carbon fixation, fatty acid biosynthesis, carotenoid biosynthesis, isoprenoid biosynthesis, and chlorophyll biosynthesis. Colour codes: green, nucleus encoded, present in predicted chloroplast proteome; amber, plastid encoded, present in predicted chloroplast proteome; light green/white, combination of green and amber in case of multiple subunits/isoforms and grey, expected but not found.

While it was initially thought that organisms possessing plastids of red versus green algal origins would have exclusively or overwhelmingly genes derived from transfer of endosymbiont genes to their nuclei, this is increasingly recognized as simplistic (Curtis et al., 2012, Dorrell et al., 2017). Since plastid acquisition in euglenoids is a relatively recent event, we sought to investigate the evolutionary origins of the genes derived from gene transfer. While we identified a set of genes best explained by gene transfer from the mitochondria, we also identified a notable cohort of over 1000 genes representing probable nuclear transfer of endosymbiotic genes as a result of acquisition of the plastid. These orthogroups are enriched for regulatory functions, as well as nucleic acid binding and oxidoreductase GO terms. Significantly, the data suggests complex acquisition in terms of gene complement, with contributions from green, red and brown plastids, consistent with earlier work based on a much less extensive dataset (Ahmadinejad, *et al.,* 2007, Maruyama, *et al.,* 2011). Recapitulating the Orthofinder results, the analysis from single protein phylogenies described above found that the largest bin of phylogenies (572 proteins or 14%) not showing a vertical pattern of descent related *E. gracilis* proteins to those from green plants and algae, likely representing genes acquired by the endosymbiotic gene transfer from the ancestor of *Euglena* secondary chloroplast. This category is followed by four bins of genes related to four algal groups – haptophytes, cryptophytes, ochrophytes and chlorarachniophytes. While many proteins within the chlorarachniophyte bin may represent mis-assigned genes related to green algae, the relatively large numbers related to the three brown-algal groups (723 in total) suggest that these algae contributed considerably to the *Euglena* genome and that the process of chloroplast endosymbiosis might be more complex. On the other hand, the number of proteins related to red algae and glaucophytes (50 and 53) seems negligible. The contributions of multiple algal lineages could be explained by phylogenetic sampling issues but are also consistent with the "shopping bag hypothesis” for plastid origins, that suggests an extended process of transient endosymbiosis and gene acquisition by the host prior to permanent plastid establishment (Howe et al., 2008).

Protein targeting to the secondary plastid in *Euglena* involves trafficking *via* the Golgi body. Since the organelle was newly established in the euglenoid lineage, this implies that at least two novel membrane-trafficking pathways should exist in *Euglena* compared with other excavates, one anterograde *trans*-Golgi to plastid and a retrograde plastid to *trans*-Golgi pathway. The relevant machinery for such pathways could be produced *via* either gene transfer from the green algal host or duplication of host membrane-trafficking machinery. We found no reliable evidence for contributions to the endomembrane protein complement by endosymbiotic gene transfer, but did find extensive gene duplications within the endomembrane machinery (Figure S7). Specifically, additional paralogues of key factors involved in post-Golgi to endosome transport e.g. AP1 and Rab14 are present, as are expansions in retromer and syntaxin16 that specifically serve to retrieve material from endosomes to the *trans*-Golgi network.

Overall, these data are consistent with a shopping bag origin, i.e. a period of kleptoplasty, sampling plastids from various algal origins, prior to the stable establishment of the secondary green plastid, as well as the model that novel pathways are established by gene duplication and co-evolution of interacting organelle identity factors, as described by the Organelle Paralogy Hypothesis (Dacks and Field, 2007).

#### Potential for biotechnological exploitation

Finally, we investigated genes relevant to features of *Euglena* that make this organism of biotechnological interest. These include aerobic and anaerobic capacity, unusual tolerance to heavy metals, lipid biosynthetic ability and possible genetic manipulation as well as other pathways (see supplementary information).

*Euglena* appears to rely on a restricted, if highly unusual set of pathways for anaerobic ATP production. In contrast to anaerobic eukaryotes and trypanosomatids *Euglena* lacks capacity to couple production of acetate as a metabolic end-product with ATP production through substrate-level phosphorylation, as well as respiration *via* exogenous electron acceptors other than O_2_. Lactate and succinate are predicted to be the only major end-products of anaerobic catabolism, as suggested previously (Tomita et al. 2016). Additions to the known (Zimorski et al. 2017) anaerobic repertoire of *E. gracilis*, however, include a pyruvate:formate lyase (PFL) plus its activating enzyme and a gene encoding a flavodiiron protein. PFL appears as a common trait for chlorophyte green algae and protists possessing chloroplasts of green algal origin (Ginger et al. 2010), while the flavodiiron protein is found in anaerobic protists, heterotrophs and chlorophyte algae. In *Euglena* this system may provide defence against reactive oxygen species in either the chloroplast and/or anaerobic conditions. PFL provides another route for conversion of pyruvate to acetyl-CoA and is utilised by *Chlamydomonas* in both the chloroplast and mitochondrion under anaerobic conditions (Atteia *et al.* 2006). In summary, *E. gracilis*, known for adaptability to hypoxia and anoxia, utilizes pathways for anaerobic ATP production that are different to the pathways seen in similarly metabolically flexible eukaryotes, including the heterolobosean *N. gruberi* and the green algae *Chlamydomonas* and *Volvox carteri* or obligate anaerobic protists such as *Giardia*, *Trichomonas* and *Entamoeba.* This provides an intriguing contrast with potential diverse signaling mechanisms such as the adenylate cyclase family linked to PAS, BLUF and other sensory perception-associated domains present in both *Euglena* and *Naegleria*, emphasizing adaptability towards variable environmental conditions.

In line with the potential for *E. gracilis* to adapt to varied and harsh conditions, including anthropogenically-impacted environments, we also considered genes that may facilitate this capacity. There is a very small repertoire of Ras-like GTPases, with only two Ras orthologs present, together with two Gtr/Rag GTPases that likely interact with the TORC system and suggesting that signaling complexity and responsiveness to environmental challenge may reside elsewhere. There is a moderate to very large family of the ABC, P-type ATPases and SLC transporters. For instance, 5 ABC, 5 P-type ATPases, and 39 SLC belonging to the ATP binding cassette and SLC superfamilies were predicted, with multiple copies in some instances, suggesting active uptake of nutrients and complex mechanisms in resilience to environmental insult (Table S5).

Pathways for amino acid biosynthesis absent from parasitic trypanosomatids, *B. saltans* and *N. gruberi* are present in *E. gracilis* (Table S5). There are multiple fatty acid biosynthetic pathways in Euglena (Zimorski et al., 2017). Intriguingly, a single predicted polypeptide of 4121 amino acids, encoding all the necessary subunits of a multifunctional fatty acid synthase (EG_transcript_9) correlates well with early isolations of a huge, cytosolic multi-functional fatty acid synthase from *Euglena* (Walker et al. 1981 inter alia). Genome-led annotation of the 4121 type I FAS contrasts with the textbook descriptions of type I FAS enzymes in animals and yeast.

The other major class of lipids required by most, if not all, eukaryotes are isoprenoids. Like in other phototrophs, a MEP/DOXP pathway, presumably plastidic, is present, but a single HMG-CoA reductase (enzyme of the mevalonate pathway) is found in *Euglena* of the same length as the trypanosomatid enzyme and lacking N-terminal TMDs that characterise animal, yeast, and plant isoforms, and raises the possibility of a mitochondrial localization, hitherto seen only in trypanosomatids and which contrasts with the textbook descriptions of ER and peroxisomal localized HMG-CoA reductases in animals, yeast, and plants. In trypanosomatids acetate formed as an end-product in the mitochondrion is transported into the cytosol for lipid biosynthesis. In the absence of ATP-dependent citrate lyase or enzymes likely to produce acetate from mitochondrial acetyl-CoA from *Euglena* genome and transcriptomes reported here, *Euglena* sources acetate or acetyl-CoA for cytosolic biosynthesis of fatty acids or other compounds for this second isoprenoid synthesis pathway via alcohol and aldehyde dehydrogenases..

In summary, the data presented here offers a huge number of gene sequences for manipulation, relating to potentially important biotechnologically relevant pathways for the first time. Promisingly, there are sufficiently complete sets of both the meiotic and RNAi machinery encoded in the genome (Table S5) to suggest that forward genetics and RNAi knock-down are possible (Tomiyama et al., 2017), facilitating the development of Euglena into a model cell biological system and biotechnological tool.

### Conclusions

*Euglena gracilis* has experienced a resurgence in interest recently, for its potential applications in biotechnology, its basic biology and its evolutionary history. We present here a thorough analysis of the coding complement of *E. gracilis*, together with insights into the organisation of the genome. The genome of *E. gracilis* is extremely large, consistent with many earlier estimates (Ebenezer *et al.,* 2017), highly repetitive and has exceptionally low coding content. The remarkable cell biology of euglenids has been known from early morphological studies, and includes a striated surface and photosensing eyespot, the latter a lineage-defining feature.

We predict a highly divergent surface proteome, and expansions of potential signaling proteins that are also likely present at the plasma membrane. *Euglena* possesses the machinery for synthesis of lipophosphoglycan, which it shares with *Phytomonas* and *Leishmania*, and which suggests these systems may have evolved as defensive system subsequently subverted to roles in pathogenesis (Eggimann, *et al.,* 2015). Significantly, we find evidence for a pathway for evolution of euglenid nuclear systems, with gradual loss of conventional kinetochores, cis-splicing and complex RNA processing at the NPC.

Rather unexpectedly, there is little evidence for transcriptional control, which is highly similar to kinetoplastids. Significant reliance on post-transcriptional processes has been long recognized as a feature of *Euglena* (Saint-Guily et al., 1994), but there is also mounting evidence that post-transcriptional, translational and degradation regulation are crucial determinants of protein abundance (Vogel and Marcotte, 2012).

An extensive endomembrane system indicates complex internal organization and multiple endosomal routes representing mechanisms for the sorting, uptake and digestion of material from a range of sources. We also find evidence for novel trafficking pathways between the endomembrane system and the chloroplast; this, together with analysis of the plastid proteome, provides insights into the processes by which secondary plastids become enslaved. Experimental analysis of the plastid will be required to validate these reconstructions, and is currently underway.

In summary, we describe here a ‘complete’ ORF dataset for *E. gracilis*, which is suitable for molecular level analysis, proteomics and pathway reconstruction. We hope that these data will usher in a new phase of analysis and utilisation of this fascinating organism, and permit a full appreciation of Euglena and its contributions to the ecosphere.

### Materials and methods

#### *Cultivation of* E. gracilis

*E. gracilis* strain Z1 was the kind gift of William Martin (Heinrich Heine University, Düsseldorf). Cells were cultivated at room temperature under continuous illumination from a 60 watt tungsten filament bulb at 20 cm from the culture vessel in Hutner‘s media (Hutner, 1966). Cells were collected at exponential phase of growth at ~9 x 10^5^ cells/ml, with cell density measured using a haemocytometer. For light and dark adaptation, cells were adapted to Hutner heterotrophic medium (Hutner, 1966) for 16 days prior to the initiation of a light or dark growth period. Cultures were subcultured and dark-adapted cultures were transferred to a light proof box adjacent to the light cultures. Subculturing was done under low light conditions periodically and cultures were maintained for up to two weeks prior to harvesting. The impact of a prolonged period under dark conditions was assessed by microscopy (Zeiss LSM 700 confocal microscope; x40 Plan-Neofuar NA1.3 lens under phase contrast and by UV/VIS spectroscopy using a Shimadzu UV-2450, wavelength scan of 190 nm – 800 nm.

#### Isolation of RNA and protein for gene expression studies

Equivalent numbers (1 x 10^7^ cells) of dark or light cultured cells were harvested by centrifugation at 25°C, 1000 xg for 10 mins. RNA extraction was performed using the Qiagen RNeasy Mini Kit (Cat. No. 74104). Genomic DNA contamination was eliminated by performing an on-column DNase digestion step. Extracted RNA was preserved in −80°C for RNA sequencing. For proteomics, cells were washed with PBS containing Roche complete protease inhibitors, extracted with standard NuPAGE LDS sample buffer (3X), sonicated and lysates containing 1 x 10^7^ cells were fractionated on a NuPAGE Bis-Tris 4-12% gradient polyacrylamide gel under reducing conditions. The sample lane was divided into eight slices that were then subjected to tryptic digest and reductive alkylation.

#### Proteomics analysis for gene expression studies

Liquid chromatography tandem mass spectrometry (LC-MS^2^) was performed in house at the University of Dundee, UK. Samples were analyzed on an Dionex UltiMate 3000 RSLCnano System (Thermo Scientific, Waltham, MA, USA) coupled to an Orbitrap Q-exactive mass spectrometer (Thermo Scientific) at the University of Dundee FingerPrints Proteomics facility and mass spectra were analyzed using MaxQuant version 1.5 (Cox and Mann, 2008) searching the predicted *E. gracilis* proteome (*de novo* transcriptome assembly) reported here. Minimum peptide length was set at six amino acids, isoleucine and leucine were considered indistinguishable and false discovery rates (FDR) of 0.01 were calculated at the levels of peptides, proteins and modification sites based on the number of hits against the reversed sequence database. Ratios were calculated from label-free quantification intensities using only peptides that could be uniquely mapped to a given protein. If the identified peptide sequence set of one protein contained the peptide set of another protein, these two proteins were assigned to the same protein group. P values were calculated applying t-test based statistics using Perseus (Tyanova et al., 2016). 8661 distinct protein groups were identified in the MaxQuant analysis. For further analyses data were reduced to 4297 protein groups by rejecting those groups not identified at the peptide level in each of the three replicates for one state. Additionally, a cohort of 384 protein groups was extracted that were observed in only one state (232 light and 152 dark).

#### Ultrastructure of *E. gracilis* cells in light and dark conditions

Two populations of *Euglena gracilis* cells cultured in either light or dark conditions were initially fixed using 2.5% glutaraldehyde and 2% paraformaldehyde in 0.1M sodium cacodylate buffer pH7.2. Both samples were post-fixed for1 hour in buffered 1% OsO_4_ and embedded in molten agarose prior to incubating overnight in 2% uranyl acetate. Agarose pellets were dehydrated through a graded acetone series and slowly embedded in Low Viscosity resin (TAAB Ltd.) over 4 days. Following polymerization, 70-90nm thin sections were cut by ultramicrotome, post-stained using 2% uranyl acetate and Reynolds lead citrate and imaged with a Hitachi H-7650 transmission electron microscope. Image resolution varied between 20 and 0.3nm per pixel depending on magnification.

#### Transcriptome analysis for gene expression studies

Extracted RNA was sequenced at the Beijing Genomics Institute (http://www.genomics.cn/en/index). Analysis and comparisons of the data were performed using standard pipelines. An estimated 62M clean reads were generated which were subject to quality filtering using Trimmomatic (Bolger, *et al.,* 2014), to remove low quality bases and read-pairs as well as contaminating adaptor sequences prior to assembly. Sequences were searched for all common Illumina adaptors and the settings used for read processing by Trimmomatic were LEADING:10 TRAILING:10 SLIDING WINDOW:5:15 MIN LEN: 50. The trimmed filtered reads were then used to quantify the *de novo* assembled transcriptome using Salmon (Patro et al., 2016) with the bias Correct option. Expected counts were integerised before being subject to differential expression testing using DESeq2 (Love et al., 2014) using default parameters. In the transcriptomics analysis, 66542 distinct sequence classes were detected and the data was reduced to 41045 applying the same rejection criteria as the proteome (minimum three replicates).

#### Nucleic acid isolation and purification for genomic and transcriptomic studies

*E. gracilis* genomic DNA was isolated using the Qiagen DNA purification system to obtain low and high molecular weight DNA for Illumina paired-end and mate-pair read libraries (100bp paired-end libraries with insert sizes of 170bp, 500bp, 800bp, and mate-pair libraries with insert sizes of 2kbp, 5kbp and 40kbp). For the shorter length libraries (≤ 5kbp), cells were harvested by centrifugation for 10 mins at 1000 g, and DNA was extracted using the Qiagen DNAeasy blood and tissue kit (Catalogue No. 69504). The procedure designed for cultured animal cells was modified and involved firstly, using 1 x 10^7^ cells and secondly, prior to adding Buffer AL, 200 μl of RNase A was added to sufficiently eliminate RNA contamination. Immediately after the washing step with Buffer AW2, centrifugation was performed for 1 min at 20,000 g to eliminate traces of ethanol. To obtain high molecular weight DNA fragments for the ≥ 40kb insert size library, the Qiagen Genomic-DNA isolation kit (blood and cell culture DNA kit - Maxi, Catalogue No. 13362) was used. In this case 1 x 10^8^ cells were harvested. Prior to adding Buffer C1, samples were ground in liquid nitrogen using aplanetary ball mill at 300 rpm for 3 min (the grinding was limited to two cycles to minimise shearing of DNA). Four wash steps were performed to remove contaminants including traces of RNA. To determine molecular weight, 400 ng of DNA was loaded onto a 0.45 % agarose gel in TAE buffer/ethidium bromide system, stained with Thermo Scientific 6X Orange Loading Dye, and electrophoresed at 80 V for 2 hrs. A NanoDrop spectrophotometer (DeNovix DS-11+) was used to determine concentration and purity. Total RNA from *E. gracilis* was isolated using the Qiagen RNeasy Mini kit (Cat. No. 74104) and the protocol for the purification of total RNA from animal cells using spin technology was employed as above.

#### Library preparation and sequencing for genomic and transcriptomic studies

Genome and transcriptome library preparation and sequencing were performed at the Beijing Genomic Institute, using Illumina Genome Analyzer HiSeq 2000 and MiSeq. In the former case, paired-end genomic sequence of multiple read lengths (49 bp and 100 bp) corresponding to eight insert size libraries (170 bp, 250 bp, 500 bp, 540 bp, 800 bp, 2kbp, 5 kbp, and 40 kbp) were generated with a combined length of ~ 57 Gbp. Additional PacBio libraries were generated at the University of Washington) (5.5 Gbp combined length) and Université Paris-Sud (3.3 Gbp combined length) and the data were kind gifts. A combined total of 914,547 PacBio sequenced reads were generated with estimated average length of 9560.59 and an estimated coverage of 4.4X respectively.

#### Genome and transcriptome assembly

Multiple routes were explored for the generation of an acceptable assembly (Ebenezer et al., 2017). The most successful strategy, as assessed by Core Eukaryotic Gene Mapping Analysis (CEGMA) and the proportion of RNAseq reads that mapped to the genome assembly (Hornett and Wheat, 2012; O’Neil and Emrich, 2013) utilised Platanus, SSPACE and SGA. Assembling the quality controlled error corrected reads with Platanus, followed by subsequent rounds of scaffolding with SSPACE and gap filling using SGA, resulted in a *de novo* assembly with an N_50_ of 955bp, comprising 2,066,288 scaffolds (Table S1). Of note was that 87% of RNAseq reads from the transcriptome successfully mapped to this assembly. A transcriptome was also assembled using a combination of data generously provided by Rob Field (O’Neill, *et al.,* 2015) and additional data generated in house (Table S1). Reads were quality filtered using Trimmomatic and searched for Illumina adaptors. Ribosomal RNA sequence was removed using SortMeRNA using default settings, before read error correction using Bayes

Hammer with default settings. Reads were normalized using khmer with settings –C 20 -k 21 -M 8e9 and overlapping paired-end reads joined using ALLPATHS-LG and all reads subject to *de novo* assembly using SGA, minimum overlap size of 80 nucleotides, no mismatches. These assembled contigs were reassembled using SGA, with each round reducing overlap size by 10 nucleotides. The filtered, normalized joined reads were then mapped to this assembly using Bowtie2. Reads that were absent from the assembly were identified and combined to the assembly file. The combined un-assembled reads and assembled contigs were subject to assembly using SGA with an overlap size of 70. This process of identifying unmapped reads and reassembling with SGA was repeated each time decreasing the overlap size by 10 nucleotides. Contigs were then subject to scaffolding using SSPACE and the full set of non-ribosomal, corrected, normalized paired-end reads using the settings -k 10 -a 0.7 -n 50 -o 20. Scaffolds were subject to gap filling using the SGA gap filling function. Finally, the assembled contigs were subject to base-error correctin using Pilon with the default settings. CEGMA (Parra and Bradnam, 2007) suggests ~88% completeness in terms of representation of coding sequence.

#### Genome and transcriptome structural and functional automatic annotation

*In silico* analysis such as open reading frame (ORF) determination, gene predictions, gene ontology (GO) and KEGG (biological pathways) and taxa distribution were performed as part of an automatic functional annotation previously described by Kaneshisa, *et al.,*2004, 2010, with minor modifications. Six frame translation and ORF determination of assembled transcriptome sequences were predicted using TransDecoder prediction tool (https://transdecoder.github.io/) and GMST, and the longest ORF with coding characteristics, BLAST homology, and PFAM domain information extracted (Stein, *et al.,* 2002). The predicted ORF was queried against the NCBI non-redundant protein database using BLASTp homology searches, and the top hit for each protein with an E-value cutoff >1e^-10^ retained. Using the Blast2Go automatic functional annotation tool (Conesa, *et al.,* 2005), the GO annotations of the best BLAST results with an E-value cutoff >1e^-10^ were generated from the GO database. The protein domain, biological pathway analyses, and top species distributions were determined using InterPro, BLAST, enzyme code and KEGG. To greatly reduce run times, BLASTp and Interpro scans were processed locally prior to uploading to Blast_2_Go in .xml file formats.

#### Assembling sequence data, data mining and phylogenetic inference

Homology searches for orthologs and paralogs of specific biological annotations were performed against the predicted proteome for *E. gracilis* using BLASTp. Clustering at 100% identity was performed for the predicted *E. gracilis* proteins using the Cluster Database at High Identity (CD-HIT) algorithm (Li and Godzik, 2006) to remove gapped/incomplete and redundant sequences. Sequences with significant BLASTp top hit search (E value = 1e^-10^) were subjected to both Reversed Position Specific Blast *RPS*-*BLAST* (NCBI CDD Search) and InterProScan. The annotated sequences with domain and/or protein signature matches were extracted using a combination of custom UNIX commands and BioPerl scripts, and clustered to 99% identity using CD-HIT. CD-HIT outputs a set of 'non-redundant' (nr) protein representative sequences which were aligned to known eukaryotic protein reference sequences using the Multiple Sequence Alignment (MSA) tool, ClustalX2 (Larkin, *et al.,* 2007) and MAFFT (Katoh and Standley, 2013). Poorly aligned positions or gaps were removed using the gap deletion command prior to alignment, and the final alignments processed locally for phylogenetic inference with the PhyML Command Line Interface (CLI) using default settings (Guindon and Gascuel, 2003; Le and Gascuel, 2008; Yang, 1993; Guindon *et al.,* 2010), RaxML (Stamatakis, 2012), FastTree (Price, *et al.,* 2010) and MrBayes (Huelsenbeck and Ronquist, 2001). Annotations of the trees were performed using TreeGraph2 (Stöver and Müller, 2010) and Adobe Illustrator. For analytical methods of specific aspects of the biology of *E. gracilis*, which include, amongst others, predicted surface protein cohort, mitochondria and the plastid proteomes see supplementary data.

#### Contigs (>10kbp) in the *E. gracilis* genome

For an initial insight into the architecture of the genome contigs >10kbp were analysed. These contigs were interrogated using tblastn with the *E. gracilis* proteome predicted from the transcriptome. Sequences with hits were further interrogated using the Exonerate algorithm for insights into splicing mechanisms and coding regions using the --protein2genome and -showquerygff and --showtargetgff options. Sequences, and their respective splicing coordinates in gff3, were uploaded to the Artemis genome viewer for visualization. Coding regions in gff formats were extracted and translated using a combination of bedtools (getfasta) and the EMBOSS getorf tools.

#### Orthologous group clustering

To identify orthologous genes in *E. gracilis* shared across eukaryotic taxa, we clustered the *E. Gracilis* predicted proteome with thirty selected eukaryotic taxa using Orthofinder (Emms and Kelly, 2015) with taxa distribution including: kinetoplastids, other members of the excavates, unikonts, bikonts, green algae, land plants, red algae, and secondary host.

*Phylogenetic analyses of ancestry of* Euglena *genes.* All 36,526 predicted nucleus-encoded proteins were blasted (blastp 2.2.29) against a custom database containing 208 organisms (Table 2C). Homologues with e-value < 10^-2^ were retrieved. Since an unrooted phylogenetic tree can be calculated only for < 3 organisms, all proteins with less than three recovered homologues (16,636 proteins) were excluded. The remaining (19,890 proteins) were aligned (MAFFT 7.273; default parameters) and trimmed (Trimal 1.2, default parameters). Alignments longer than 74 amino acid residues and with all sequences determined, i.e. there was no sequence containing only undetermined characters, (18,108 alignments) were used for tree reconstruction. The trees were calculated with RAxML (v8.1.17; 100 rapid bootstraps) in Metacentrum (The National Grid Infrastructure in the Czech Republic). Custom scripts (python 3.4) was used to sort the trees into bins based on the taxonomic affiliation of the sister branch of *Euglena*. The tree was included in a bin if one of the two *Euglena* sister branches in the unrooted tree contained only taxa from one of the defined taxonomic groups and the sisterhood was supported by bootstrap 75 % or higher (Table 2C). In 34 cases, in which both sister branches contained taxa from one defined group, but each branch different group, the tree was assigned to the two respective bins.

#### Mitochondrial proteome prediction

The predicted proteins were subjected toBlast2GOtoBlast2GO (https://www.blast2go.com/) and KAAS (Moriya, *et al.,* 2007) automatic annotation, BLASTp searches against the *T. brucei*, *Homo sapiens*, *Saccharomyces cerevisiae*, and *Arabidopsis thaliana* reference mitoproteomes and, finally, targeting signal prediction usingTargetPusingTargetP (Emanuelsson, *et al.,* 2000). *E. gracilis* protein was predicted as mitochondrial if: i) its TargetP mitochondrial score was higher than 0.9 (607 proteins), or ii) there was an ortholog in at least one reference mitoproteome, not associated with non-mitochondrial functions (343 proteins), or iii) it was assigned mitochondrial by Blast2GO (with the exception of the MTERF family) (62 proteins).The missing members of the found mitochondrial pathways and modules were identified by a manual search (81 proteins). To streamline the final annotated output, and to ensure retention of only the most reliable predictions, we chose the most confident annotation between Blast2GO, BLASTp and KAAS for each protein. The final mitochondrial dataset includes1093 proteins.

#### Plastid proteome prediction

The translated *E. gracilis* transcriptome (predicted proteome) was subjected to signal prediction pipeline using a combination of SignalP (Petersen et al., 2011) and PrediSI (Hiller, *et al.,* 2004) while chloroplast transit peptide prediction was performed using ChloroP (Emanuelsson, *et al.,* 1999). The sequences which scored positive by either SignalP (2551 sequences) or PrediSI (4857 sequences) were cut at the predicted signal peptide cleavage site. The sequences were then truncated to maximum length of 200 amino acid residues for faster calculation and analyzed by ChloroP. The preliminary dataset of *E. gracilis* plastid targeted proteins (1679 sequences) consisted of transcripts which scored positive in SignalP + ChloroP (59 sequences), PrediSI + ChloroP (1002 sequences), and SignalP + PrediSI + ChloroP (618 sequences) analysis. In the second step, model dataset of 920 sequences of *Arabidopsis thaliana* proteins localized to plastid envelope, stroma, thylakoid, grana, and lamellae obtained from the public AT_CHLORO proteomic database (Bruley, *et al.,* 2005) were searched by BLAST against the whole translated *E. gracilis* transcriptome and the identified homologs were then combined with the results of orthogroup clustering performed by OrthoFinder (see above). Based on these searches, an additional 144 sequences representing orthologs of *A. thaliana* chloroplast proteins were added to the dataset of Euglena predicted plastid proteome regardless of their targeting sequences. This enriched dataset of 1823 proteins was annotated automatically using BLAST at NCBI, KOBAS (Xie, *et al.,* 2011) and KAAS (Moriya, *et al.,* 2007) independently. All automatic annotations including KO and EC numbers were then revised and edited or corrected manually and used for metabolic map reconstruction. The missing enzymes and subunits of otherwise chloroplast pathways and complexes were investigated and eventually added manually to the set regardless of their targeting sequences during the manual annotation and pathway reconstruction. This approach resulted in inclusion of another 79 sequences. The final set of predicted *E. gracilis* chloroplast proteins consisted of 1902 entries.

## Acknowledgements

This work was supported by the Yousef Jameel Academic Program (through the Yousef Jameel PhD Scholarship), the Cambridge Commonwealth, European and International Trust, the Cambridge University Student Registry, the Cambridge Philosophical Society (all to TEE), the Medical Research Council (Grant #: P009018/1 to MCF), and German Aerospace Center – DLR, Cologne, on the behalf of Federal Ministry of Education and Research (BMBF), Germany (Grant no: 50WB1128 and 50WB1528 to ML), the European Research Council CZ LL1601 and the Czech Grant Agency No. 16-21974S (to JL), Czech Ministry of Education, Youth and Sports - National Sustainability Program II (Project BIOCEV-FAR) LQ 1604 and by the project ‘‘BIOCEV’’ (CZ.1.05/1.1.00/02.0109) and by the Czech Science Foundation project nr. 16-25280S (to VH, AV and PS). MCF is a Wellcome Trust Investigator and SK is a Royal Society University Research Fellow. Work in SKs lab was supported by the European Union’s Horizon 2020 research and innovation programme under grant agreement number 637765. We are most grateful to Purificatión Lopéz-García, David Moreira and Peter Myler for the kind donation of PacBio sequence data and also to Robert Field for permission to reutilise transcriptome data. We thank Douglas Lamont and the Fingerprints proteomics facility at the University of Dundee for excellent mass spectrometric analysis. Some computational resources were provided by the CESNET LM2015042 and the CERIT Scientific Cloud LM2015085, provided under the programme "Projects of large research, development, and innovations infrastructures”.

